# Understanding genetic diversity and phylogeography of Common Teal and its phylogenetic relationship with other water bird species in the wetlands of Kashmir Himalayas

**DOI:** 10.1101/2024.10.02.616289

**Authors:** Khursheed Ahmad, Divyanshi Bisht, Bheem Dutt Joshi, Surender P. Goyal, Parag Nigam, Khurshid Alam Khan, Iqram ul Haq, Mohsin Javid

**Affiliations:** Division of Wildlife Sciences, Sher-e-Kashmir University of Agricultural Sciences and Technology (SKUAST-Kashmir), Srinagar 190025, Kashmir, Jammu & Kashmir, India; Wildlife Institute of India, Chandrabhani, Dehra Dun, India 248001

**Keywords:** Genetic diversity, Phylogeography, Anas crecca (Common Teal), Wetland conservation, Mitochondrial DNA, Kashmir Himalayas

## Abstract

Understanding the genetic diversity and phylogeography of migratory species is critical for biodiversity conservation and the effective management of wetland ecosystems. The Kashmir Himalayas, an integral part of the Central Asian Flyway, host several key wetlands that provide critical wintering grounds for a variety of migratory birds.

This study focuses on assessing the genetic characteristics and phylogenetic relationships of the Common Teal (*Anas crecca*) in comparison to other species within the families Anatidae and Rallidae. We analysed 149 blood samples, including 71 from *A. crecca* and 78 from other species in the two families, collected from wetlands in the Kashmir region. Using four mitochondrial markers—cytochrome oxidase subunit I (COI), cytochrome b (Cyt b), 16S rRNA, and the control region—we evaluated the genetic diversity and lineage connectivity of these species.

Our findings reveal that the mitochondrial DNA haplotypes of *A. crecca* in the Kashmir Himalayas are shared with European populations, indicating strong maternal gene flow and connectivity between distant populations. A minimum spanning haplotype network analysis showed minimal nucleotide differences among haplotypes, particularly in the Cyt b and control regions, suggesting low genetic differentiation and a high degree of similarity among individuals. Notably, we identified at least four distinct maternal lineages of *A. crecca* in the Kashmir wetlands, reflecting diverse migratory sources.

Our results also highlight that DNA barcoding using COI exhibited both high and low species resolution, with significant intraspecific variation, making it a valuable tool for further phylogeographic studies. The observed genetic diversity and haplotype sharing with distant populations underscore the ecological importance of Kashmir’s wetlands as crucial habitats for migratory species. Our study emphasizes the need for targeted conservation and management strategies to preserve these vital ecosystems and the biodiversity they support.

## Introduction

The Kashmir Himalayan region is home to numerous wetlands, several of which are internationally recognized as Ramsar sites. These wetlands provide abundant resources and relatively low levels of disturbance, making them ideal wintering grounds for a wide variety of migratory bird species. Understanding the genetic diversity and population structure of these migratory birds is crucial for effective conservation and management strategies, particularly in terms of gene flow and connectivity between populations [1]. Phylogeography, which examines the geographic distribution of genetic lineages using mitochondrial DNA, has emerged as a key tool for studying these dynamics in migratory birds [2-4]. Comparative phylogeographic analyses offer valuable insights into the demographic history and intraspecific evolution of migratory species, shedding light on their adaptation to varying environmental conditions [5-9].

As migratory birds traverse different wintering grounds, understanding their genetic makeup is essential for unraveling how they adapt to diverse habitats and how different populations utilize these areas. Recent advances in DNA sequencing technology have enabled the construction of reliable phylogenies and the identification of cryptic species and population structures. The use of whole mitochondrial genomes or concatenated mitochondrial gene fragments has proven particularly powerful due to their high discriminatory power and the availability of large genetic databases [9, 10-31].

Some migratory species maintain high levels of genetic diversity by moving between breeding populations, while others exhibit weaker connectivity. Assessing genetic diversity, gene flow, and population connectivity on both global and regional scales is essential for the development of effective conservation strategies [17, 32-33]. Regional or population-specific haplotypes, which can track migration paths from multiple source populations, provide valuable insights into the migration patterns of species. However, to fully understand these patterns, harmonized genetic data across the species’ entire distribution range is needed. Mitochondrial genome characterization has proven particularly effective in phylogeographic studies, especially for widely distributed species such as ducks, geese, and swans of the family Anatidae [18, 34].

The Common Teal (*Anas crecca*), a widely distributed Anatidae species across the Holarctic, is a common wintering duck in the Kashmir Valley. It is highly abundant in all wetlands during winter, except at higher elevations in the Himalayas and the Western Ghats [35]. Although most *A. crecca* populations are observed in winter, some individuals have been recorded in the Kashmir wetlands during summer breeding seasons [36-38], and as autumn passage migrants to the Ladakh region in the Trans-Himalayas [39-41]. Outside of the breeding season, large congregations of *A. crecca* can be found in lakes, wetlands, and coastal bays, while during the breeding season, they prefer brackish and freshwater lakes and ponds in upland forests or woodlands [42]. However, populations of *A. crecca*, along with other waterfowl, face significant threats from habitat loss, poaching, pollution, and changes in land use [37, 43-45].

The wetlands of the Kashmir Valley are crucial for the wintering of a diverse range of migratory waterfowl, including the Common Teal and other species within the Anatidae and Rallidae families. Despite the ecological importance of these habitats, there is a significant lack of research on the genetic diversity, population structure, and phylogeographic patterns of these migratory birds in the Kashmir Himalayas and across India. To address this gap, our study aims to assess the genetic diversity, comparative phylogeography, and phylogenetic relationships of waterbirds inhabiting the wetlands of the Kashmir Himalayas. Specifically, we focus on the Common Teal (*A. crecca*) and its phylogenetic relationships with other species of Anatidae and Rallidae in the region.

By integrating our genetic data with global mitochondrial DNA datasets, we seek to uncover regional and global patterns of genetic diversity and lineage distribution among migratory water birds. The insights gained from this study will not only contribute to a better understanding of the genetic diversity and phylogeography of migratory birds in the Kashmir Himalayas but will also inform broader conservation efforts aimed at preserving these ecologically significant species and their habitats.

## Material and Methods

### Study Area

This study was conducted across five major wetland reserves in the Kashmir Valley: Hokersar, Haigam, Shallabugh, Wular, and Chatlum-Frushkori. These wetlands are located in the districts of Srinagar, Budgam, Ganderbal, Baramulla, and Bandipora (Figure 1), and represent critical habitats for migratory water birds.

**Figure 1.**
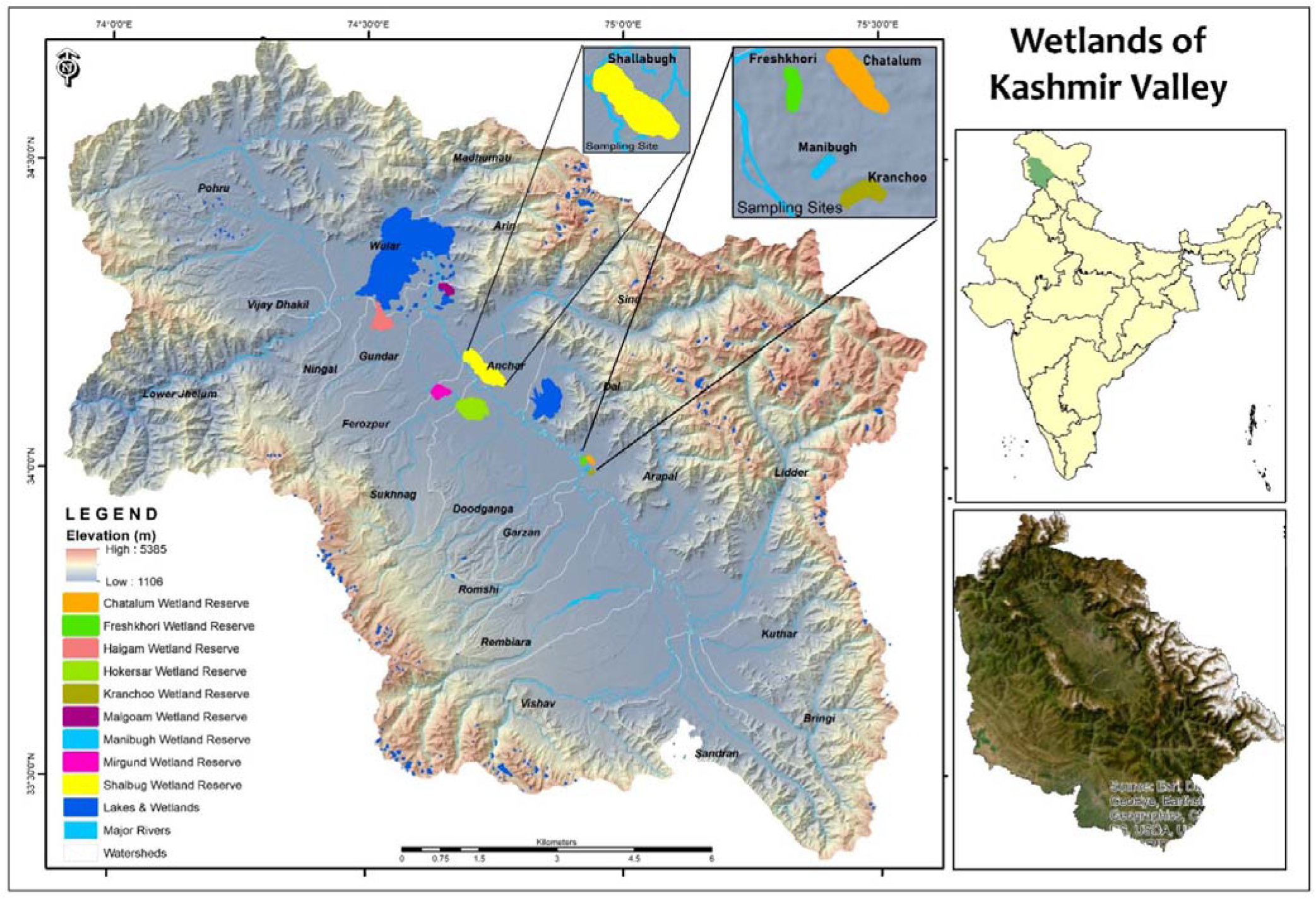
Map of the Study area indicating different wetlands of Kashmir Valley surveyed and sampled (The majority of the samples were collected from Shallabugh and Freshkuri wetlands).

### Bird Sample Collection and DNA Extraction

A total of 149 blood samples, including 71 from *Anas crecca* (Common Teal) and 78 from other water bird species, were collected from six wetlands in the Kashmir Himalayas during bird ringing and satellite PTT tagging exercises in February-March 2017 and 2018 (Table 1). The samples were stored at ambient temperature and subsequently transferred to the laboratory for DNA extraction.

**Table 1.** Details of blood samples collected from different migratory water birds from wetlands of ashmir, India during February-March 2017 & 2018.

| Common Name | Family | Species | Year of collection |  |  | Grand Total |
| --- | --- | --- | --- | --- | --- | --- |
|  |  |  | 2017 | 2018 | NA |  |
| Common Teal | Anatidae | *Anas crecca* | 64 | 7 | 0 | 71 |
| Common Moorhen | Rallidae | *Gallinula chloropus* | 14 | 8 | 0 | 22 |
| Common coot | Rallidae | *Fulica atra* | 19 | 1 | 0 | 20 |
| Northern Shoveler | Anatidae | *Spatula clypeata* | 6 | 4 | 0 | 10 |
| Gadwall | Anatidae | *Mareca strepera* | 6 | 2 | 1 | 9 |
| Mallard | Anatidae | *Anas platyrhynchos* | 6 | 2 | 0 | 8 |
| Common Pochard | Anatidae | *Aythya ferina* | 2 | 0 | 0 | 2 |
| Eurasian Wigeon | Anatidae | *Mareca penelope* | 2 | 0 | 0 | 2 |
| Graylag Goose | Anatidae | *Anser anser* | 0 | 2 | 0 | 2 |
| Tufted Duck | Anatidae | *Aythya fuligula* | 2 | 0 | 0 | 2 |
| Grand Total |  |  | 122 | 26 | 1 | 149 |

### DNA extraction

In the laboratory, bloodstains from each sample were transferred to 2 ml Eppendorf tubes. DNA was extracted using the QIAamp DNA Mini Kit (QIAGEN, Hilden, Germany) following the manufacturer’s protocol. DNA was eluted with 50 µL of elution buffer, and the extracted DNA was stored at -20°C until further analysis. Precautions, including blanks, were used to prevent contamination. DNA concentration was checked using 0.8% agarose gel electrophoresis in 1x TAE buffer, and yield was observed under a UV trans-illuminator. Samples with varied DNA concentrations were diluted to achieve a uniform concentration for downstream PCR amplification.

### PCR Amplification and DNA Sequencing

We amplified five mitochondrial gene regions—16S rRNA, 12S rRNA, cytochrome b (Cyt b), control region (CR), and cytochrome oxidase subunit I (COI)—using polymerase chain reaction (PCR). Each 10 μL reaction contained 1 μL of 1x PCR buffer, 0.5 μL of 10 mM dNTPs, 0.5 μL of 25 mM MgCl□, 0.4 μL of BSA, 0.5 U of Taq DNA Polymerase, and 2 μL of ∼20 ng genomic DNA. Both negative and positive controls were included to ensure accuracy and to monitor contamination. PCR conditions followed previously published protocols for the respective primers (Table S2). Amplification success was confirmed by visualizing PCR products on 2% agarose gels in 1x TAE buffer under UV light. Residual primers and dNTPs were removed using Exonuclease-I and Shrimp Alkaline Phosphatase (Thermo Scientific) at 37°C and 85°C for 15 minutes each. Sequencing was performed using BigDye™ Terminator v3.1 Cycle Sequencing Kit (Applied Biosystems, USA) on an ABI 3130 DNA sequencer.

### Data Analysis

Raw sequence data were edited and processed using Sequencer v.3.1 (Gene Codes Corporation, USA). Sequences were visually inspected and trimmed to uniform lengths using BioEdit v.7.0.9.0 [46]. We used NCBI BLAST to identify species from the generated sequences and validate them against reference data. Variations in sample size resulted from amplification failures or poor sequence quality. For COI sequences, the closest matches were identified using the NCBI BLAST tool, and validated sequences were processed for further analysis.

We retrieved additional mitochondrial genome sequences from NCBI for various species from different geographic locations. Multiple sequence alignments were performed using the ClustalW algorithm in MEGA X [48], and sequences were trimmed to identical lengths with no indels. Phylogenetic trees were reconstructed using the Maximum Likelihood method in MEGA X with 500 bootstrap replicates for node support. Pairwise genetic distances between haplotypes and species were calculated using the Kimura 2-parameter (K2P) distance model.

Genetic diversity indices such as nucleotide diversity (π), haplotype diversity (Hd), and mismatch distribution were computed using DnaSP v.6.12 [49]. Polymorphic sites, transversion events, and nucleotide composition (purine-pyrimidine ratio) were analyzed in MEGA X. To examine historical demographic trends, we used Bayesian skyline plots (BSP) in BEAST v.2.1.3 [51], running Markov chains for 2.5 × 10□ generations, sampling every 1000 generations, and discarding the first 2500 samples as burn-in. The substitution rate used was 2.0 × 10□□ [52], and BSP results were visualized using TRACER v.1.6 [53]

## Results

### Species Identification and Sequence Data

We identified 12 species of migratory birds with 99–100% similarity using NCBI BLAST and reference data. A total of 98 COI sequences (574 bp), 56 16S rRNA sequences (440 bp), 74 12S rRNA sequences (360 bp), 80 Cyt b sequences (370 bp), and 97 control region (CR) sequences (700 bp) were generated. All unique haplotypes identified in this study have been deposited in NCBI (16S rRNA: ON406326-ON406350; 12S rRNA: ON417475-ON417521; COI: ON387719-ON387745; D-loop: ON400515-ON400539; Cyt b: ON412323-ON412374). Further, we obtained 36 compatible sequences of *Anas crecca* from NCBI and 24 sequences from other species within the Anatidae family, using these for COI, 12S, 16S, and Cyt b to reconstruct phylogenetic relationships and evaluate genetic diversity and sequence divergence.

### Nucleotide and Genetic Diversity

The average nucleotide frequencies across the different mtDNA genes were 25.63% (A), 23.34% (T/U), 37.38% (C), and 19.93% (G) (Table S1, S2). Nucleotide composition ranged from 19.05% to 30.87% (A), 16.65% to 30.28% (T), 26.37% to 36.55% (C), and 15.69% to 23.77% (G). In the Anatidae family, the general nucleotide frequency pattern was C > A > T > G. Moreover, the GC content was relatively higher in Anatidae species.

Haplotype and nucleotide diversity varied among species. *Anas crecca*, Common Coot, Common Moorhen, and Northern Shoveler showed significant variation in both, while Gadwall and Mallard did not exhibit much variation (Table 2), suggesting stronger population connectivity across their distribution ranges. Among these species, the Common Coot had the highest haplotype diversity but low nucleotide diversity, indicating the presence of many closely related haplotypes with minimal nucleotide differences.

**Table 2.** Summary of molecular diversity of each species for different regions (concatenated) of mtDNA genome generated in the present study.

| Species name | N | nH | G+C% | S | k | h | $\pi$ | Tajima's D | Fu' FS |
| --- | --- | --- | --- | --- | --- | --- | --- | --- | --- |
| Common coot<br>(*Fulica atra*) | 18 | 7 | 0.505 | 6 | 1.392 | 0.804 | 0.0037 | -0.6650 | 7.627 |
| Common Teal<br>(*Anas crecca*) | 47 | 9 | 0.531 | 71 | 8.509 | 0.349 | 0.0231 | -1.7450 | 6.029 |
| Common moorhen<br>(*Gallinula chloropus*) | 12 | 4 | 0.500 | 4 | 0.939 | 0.455 | 0.0025 | -1.0220 | 0 |
| Gadwall (*Mareca strepera*) | 9 | 1 | 0.527 | 0 | 0 | - | 0 | 0 | 0 |
| Mallard (*Anas platyrhynchos*) | 6 | 1 | 0.535 | 0 | 0 | - | 0 | 0 | 0.351 |
| Northern shoveler<br>(*Spatula clypeata*) | 9 | 3 | 0.528 | 3 | 1.00 | 0.417 | 0.0027 | -0.3590 | 7.627 |

**N**- No of individuals; nH-Number of haplotypes; S –variable sites; k - average number of nucleotide differences; h-haplotype diversity; $\pi$ -nucleotide diversity.

The mitochondrial DNA diversity and haplotype network analysis for A. crecca indicated low genetic diversity. A minimum spanning network revealed mostly single nucleotide differences between haplotypes, except for Common Coot in the 12S rRNA, Cyt b, and control region, which suggests a recent population expansion. Although negative Tajima’s D values were observed for most species, they were not statistically significant (Table 2). The mismatch distribution analysis for A. crecca using global COI data showed a multimodal mismatch pattern, while concatenated sequences displayed a unimodal pattern (Figure 2a, 3b), indicating populations in equilibrium.

**Figure 2.**
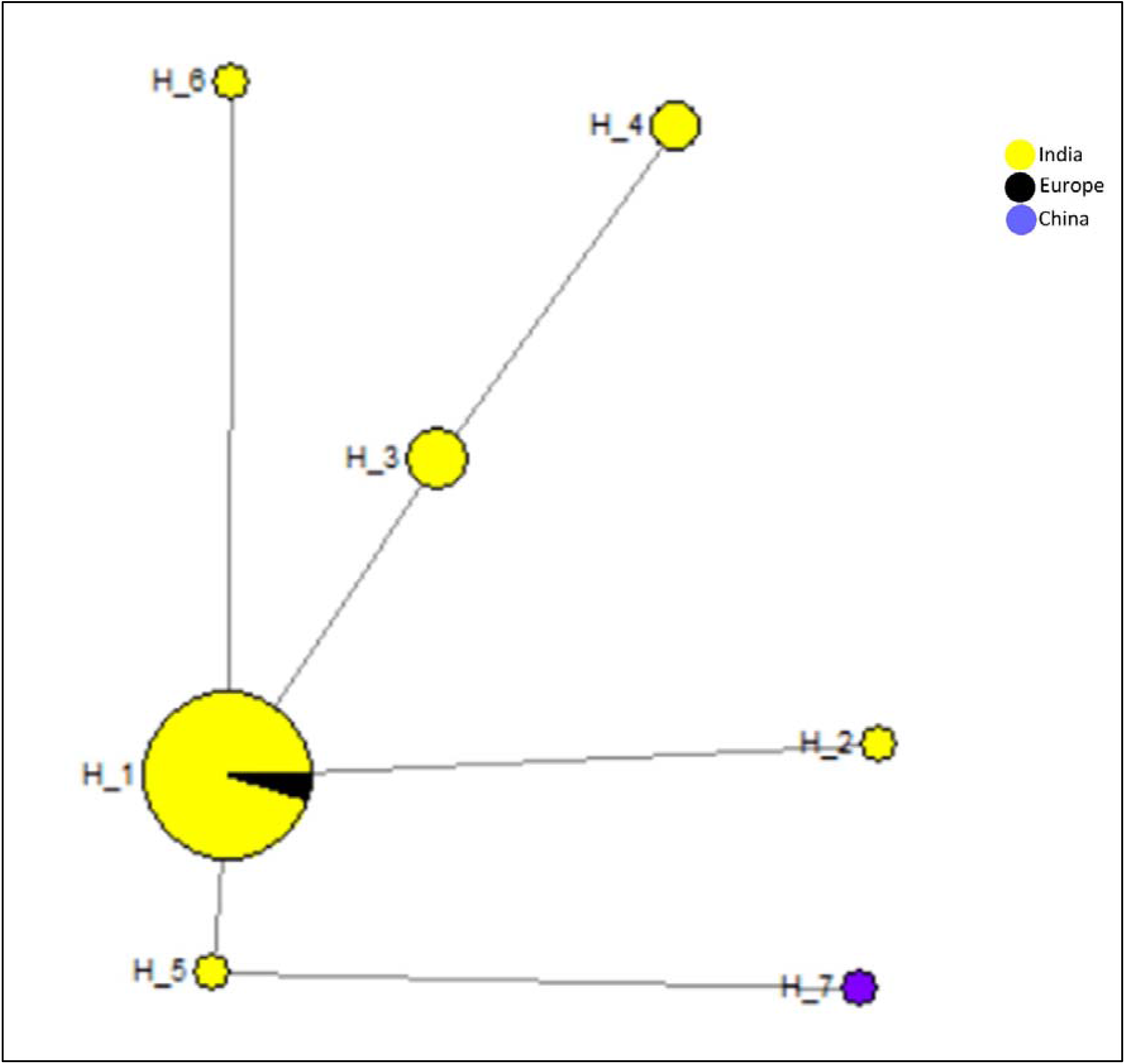
Minimum spanning network of observed haplotypes in Common Teal (*Anas crecca*) based on different mtDNA genes. Circle size is in proportional number of individuals.

**Figure 3.**
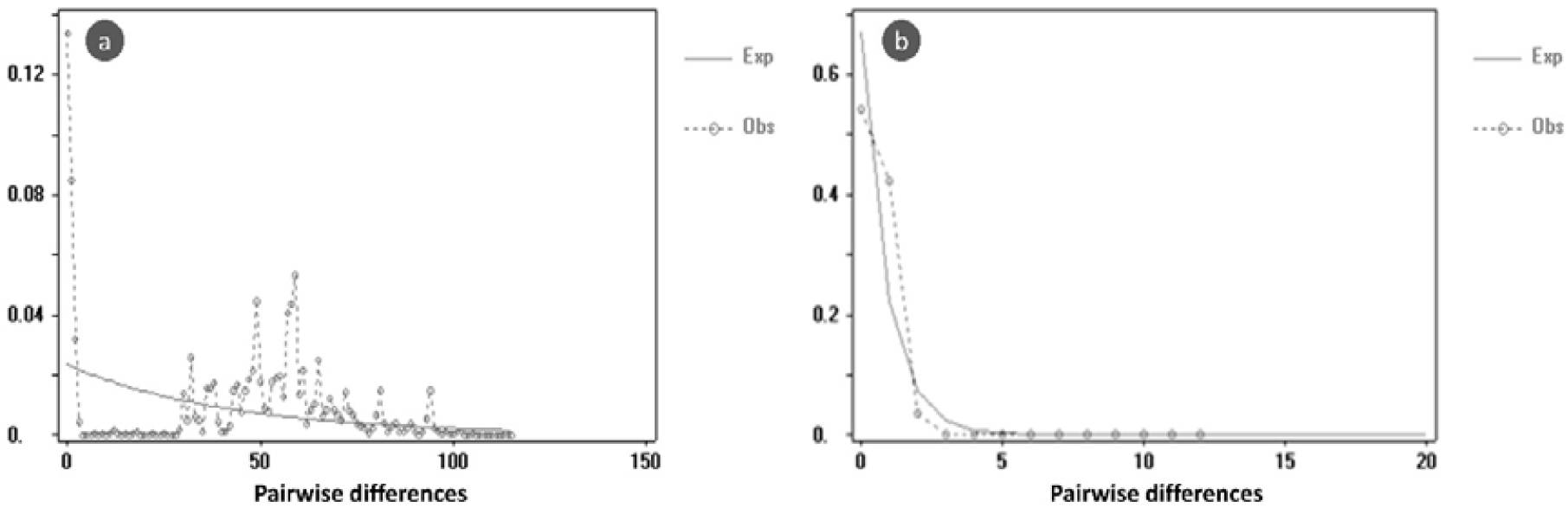
Mismatch distribution graphs of species Common Teal: a) Multimodal mismatch graph with COI; b) Unimodal mismatch graph with concatenated sequences.

Bayesian Skyline Plot (BSP) analysis for A. crecca revealed a stable population size up to 0.25 million years ago, followed by a slight decline in the effective mitochondrial population size (Figure 4). This potential decline highlights the importance of targeted conservation efforts for this species.

**Figure 4.**
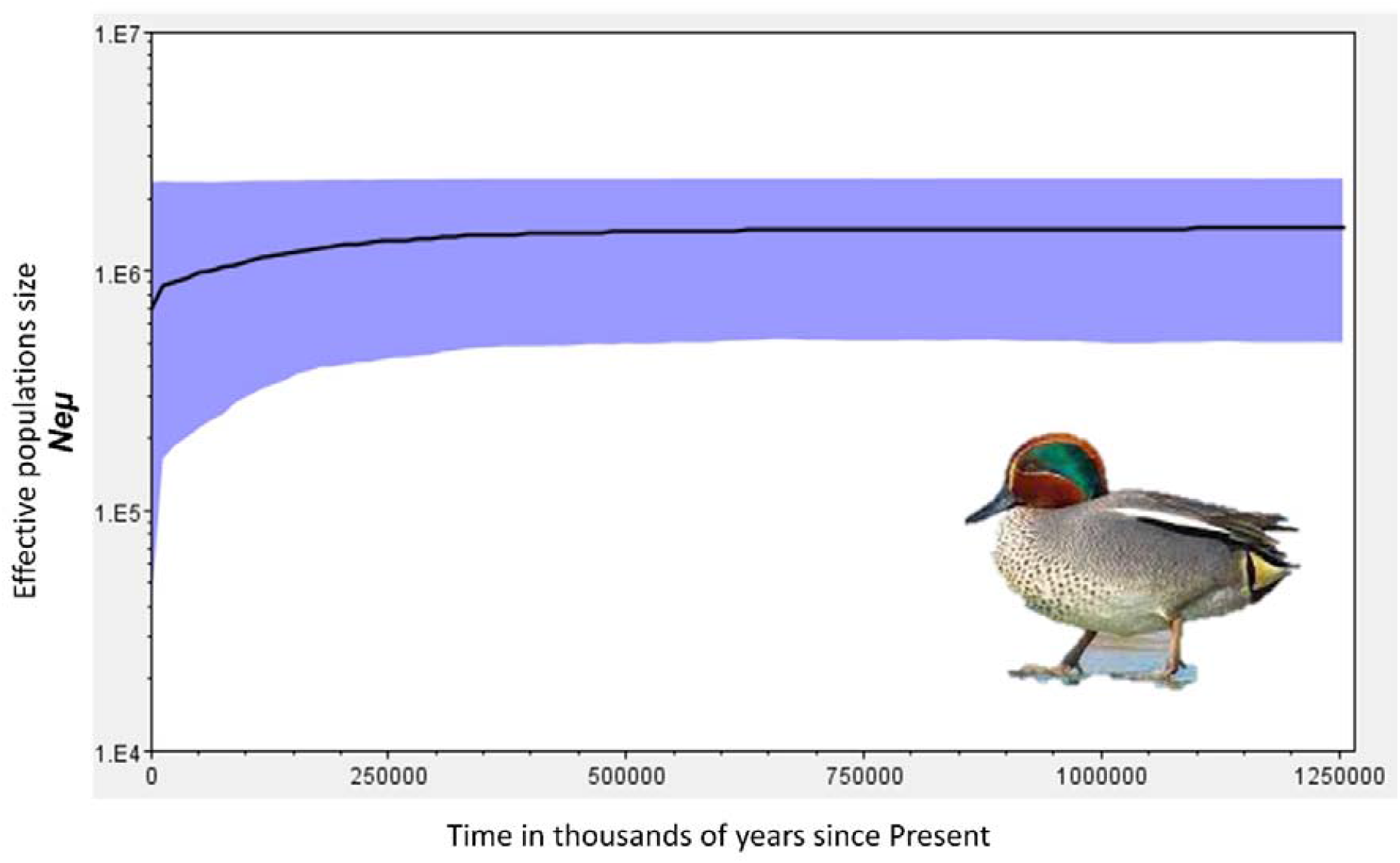
Demographic history estimated using Bayesian skyline plot (BSP) using concatenated genes of Common Teal *Anas crecca* in the major sample collection sites of Shallabugh and Freshkhori wetlands of Kashmir Valley (Ne = effective population size, μ= generation time).

### Genetic Distance and Phylogenetic Analysis

Genetic distances were calculated within and between species in the Anatidae family. The smallest interspecies genetic difference was 0.02, while the highest intraspecies distance reached 0.119. COI sequences were unique to each of the 28 bird species studied, with identical or closely related sequences found within individuals of the same species. COI sequence divergence between closely related species was lower than intraspecies variation, with some exceptions noted below.

The Neighbor-Joining (NJ) tree revealed shallow intraspecific divergences and deep interspecific divergences in most cases. The COI-based phylogenetic tree aligned well with established avian classifications, reflecting nested monophyletic lineages that corresponded with genera, families, and orders (Figure 5).

**Figure 5.**
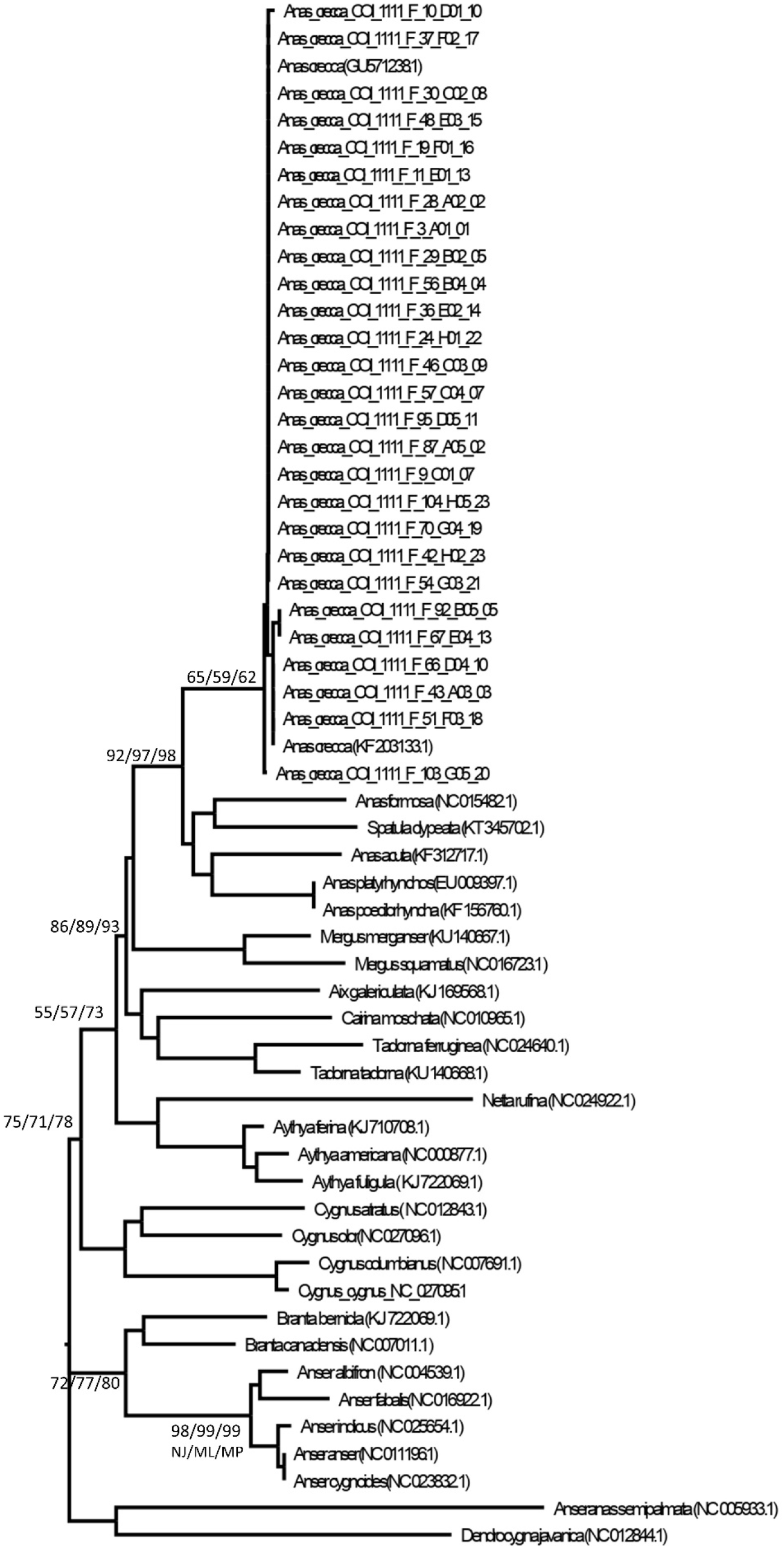
Maximum Likelihood Tree COI gene of *Anas crecca* with same family. Numbers in nodes are bootstraps values of different methods according to initial given for each methods in bottom of node. Maximum likelihood (ML), Minimum Parsimony (MP) and Neighbor Joining (NJ) method

### Phylogenetic Insights

Phylogenetic analysis across the mtDNA regions (12S rRNA, 16S rRNA, Cyt b, COI, and control region) revealed that *Anas crecca* sequences from Kashmir wetlands were shared with populations from Europe, North America, Japan, Korea, China, and Canada (Figure 5, 6). This finding underscores the strong genetic connectivity between A. crecca populations in the Kashmir Himalayas and those across its global range.

**Figure 6.**
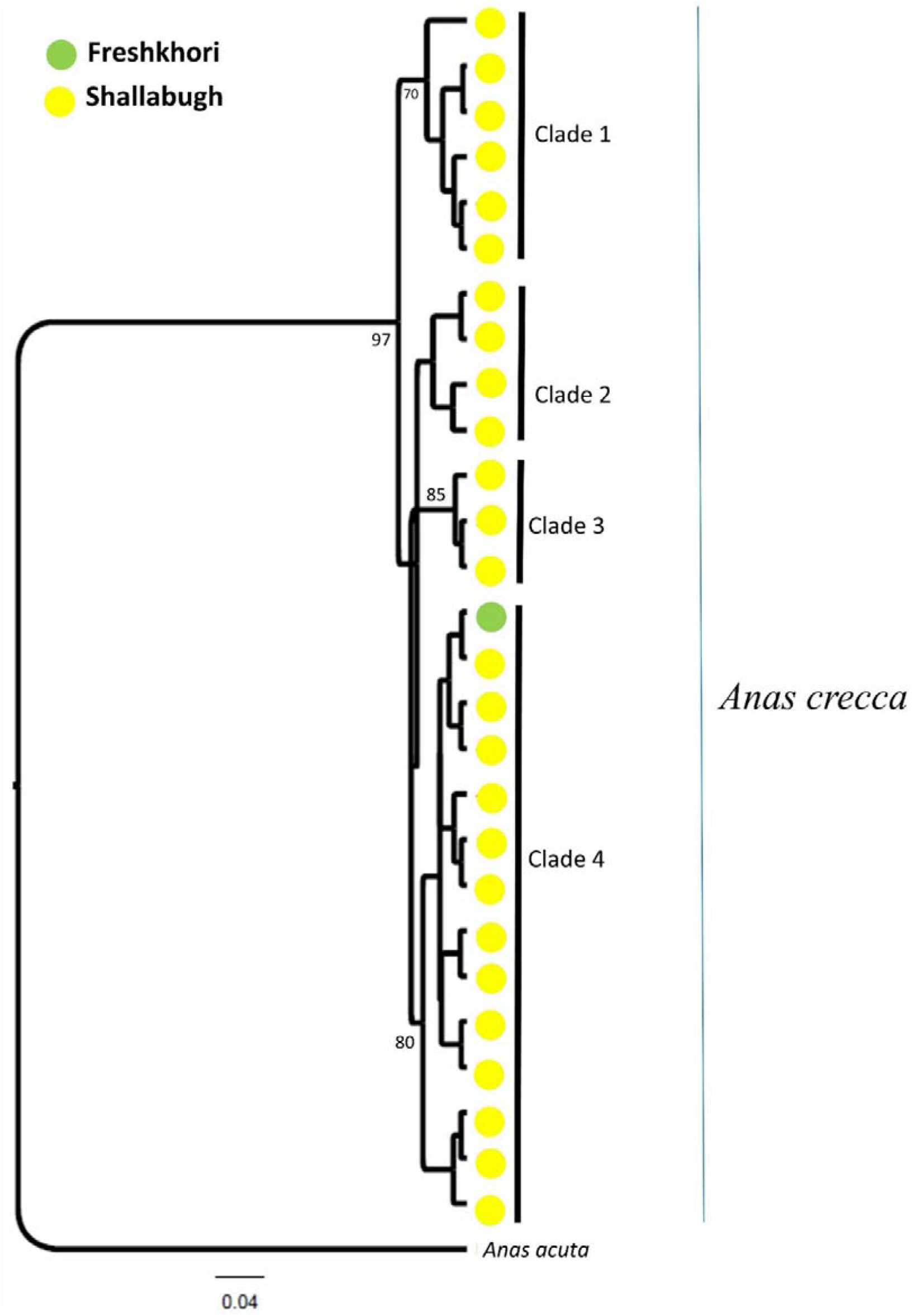
Maximum Likelihood Tree of the concatenated genes of Common Teal *Anas crecca* in the major sample collection sites of Shallabugh and Freshkhori wetlands of Kashmir Valley. Values below the node is bootstrap support and Out group species is *Anas acuta (*Accession No. KF_312717.1).

The higher haplotype and nucleotide diversity in A. crecca relative to other species suggests a larger effective population size and highlights the suitability of Kashmir’s wetlands as a critical habitat for migratory birds. Our analysis of Cyt b, control region, and COI in A. crecca further identified at least four distinct maternal lineages in Kashmir’s wetlands, suggesting migration from different geographic regions (Figure 5, 6 & Figures S1-S5).

## Discussion

Our study highlights the reduced haplotype diversity in *Anas crecca* populations, with only 9 haplotypes identified and minimal nucleotide variation (Table 2). This low genetic diversity, particularly in mitochondrial DNA, signals a potential vulnerability in the species, likely driven by a declining population size. This reduction in diversity underscores the urgent need for targeted conservation efforts, especially for *Anas crecca* populations inhabiting the fragile wetlands of the Kashmir Valley in the Indian Himalayan region.

While *Anas crecca* exhibits low genetic diversity (54), this does not appear to be a universal trend across the Anatidae family. Populations within Anatidae that have not experienced drastic declines still maintain comparable levels of genetic diversity. Genetic diversity often reflects a population’s size, ecological factors, natural history, and adaptive capacity. However, Bazin et al. (55) argue that mitochondrial DNA (mtDNA) markers may not always reflect population size or ecology. Given that Anas crecca is a migratory species, which only resides in the Kashmir Valley for limited periods, interpreting the genetic diversity based solely on mtDNA data requires caution. Additional factors such as population structure and the mixing of birds from multiple geographic sources likely contribute to the observed genetic patterns (56).

Our analysis using DNA barcoding and partitioning of genetic variation confirms that the majority of variation in *Anas crecca* occurs within populations, with only 0.10% attributed to variability among populations. This supports the findings by Hebert et al. (20, 57), who reported that nucleotide differences among individuals of the same species should typically be less than 3%.

The unimodal distribution of concatenated sequences of pairwise differences among haplotypes suggests a recent population expansion (50, 58). This is further supported by the haplotype network and mismatch distribution graphs, which indicate an excess of singleton variable sites, a common indicator of population growth (Figure 2-3). However, in contrast to global Anatidae data, the populations from our study exhibited fewer singleton variable sites. While the mismatch distribution fit well with a simulated model of population expansion, tests for neutrality (Fu’s Fs and Tajima’s D) did not support recent expansion, aligning with the haplotype network and mismatch distribution results.

If the current trend of population decline continues, the effects of genetic drift and inbreeding could further erode the genetic diversity of *Anas crecca* populations. The implications are significant: diminishing genetic diversity will reduce the adaptive capacity of these populations, making them more susceptible to environmental changes and disease. Thus, urgent conservation strategies are needed to preserve the genetic integrity of these migratory populations. Maintaining wetland habitats in the Kashmir Himalayas is critical to ensuring the survival and continued migration of *Anas crecca*.

The COI gene has proven effective as a DNA barcode in birds (57), offering a reliable tool for species identification due to its high mutation rate and capacity to differentiate between species. In this study, within-species, genus, and family K2P distances for the COI gene were found to be 0.43%, 7.93%, and 12.71%, respectively, consistent with previous studies that used COI and other mitochondrial markers for phylogenetic analysis (57, 60). Phylogenetic trees constructed using COI, along with 12S rRNA, 16S rRNA, Cyt b, and control regions, revealed congruence with existing taxonomic classifications within Anatidae (60). Yoo et al. (61) and Sun et al. (18) reported similar findings in their phylogenetic studies, with little difference observed between regional and global data. Additionally, Sun et al. (18) calculated K2P distances among 54 species within Anseriformes, showing that genetic distances between genera in Anatidae ranged from 0.037 to 0.107, while the average genetic distances were 0.103 ± 0.008. These findings were consistent with our observations, indicating significant genetic divergence between Anatidae and Dendrocygnidae, which also highlights the taxonomic robustness of COI barcoding. Furthermore, Livezey et al. (62) provided an in-depth classification of teals based on taxonomy, anatomy, and behavior, corroborating our phylogenetic conclusions.

Our findings demonstrate that *Anas crecca* in the Kashmir wetlands harbours low mtDNA diversity, a likely result of declining population sizes. This study reinforces the importance of genetic monitoring for conservation purposes and the need to protect vital wetland habitats to sustain migratory bird populations. Conservation measures must prioritize preserving the genetic diversity of *Anas crecca* and related species, thereby ensuring the ecological integrity of these populations for future generations.

## Conclusion

This study underscores the utility and limitations of DNA barcoding using COI in water bird species, revealing varying levels of species resolution. While certain species exhibited high intraspecific variation, pointing to opportunities for more detailed phylogeographic and population genetic studies, others showed a lack of divergence, particularly at higher taxonomic levels. Our comparative analysis of *Anas crecca* with other Anatidae and select Rallidae species demonstrated significant genetic divergence both within and between species. Specifically, the presence of four distinct maternal lineages in the Cyt b, control region, and COI sequences of *Anas crecca* suggests historical connectivity with populations from different regions, particularly Europe, highlighting the wetlands of Kashmir as a critical stopover for this migratory species.

Bayesian phylogenetic analysis further confirmed moderate to high divergence in *Anas crecca* and Common Coot, while species such as Common Moorhen, Northern Shoveler, Gadwall, and Mallard showed lower levels of divergence. These findings provide crucial baseline data for understanding the genetic structure of water bird species in the Kashmir Himalayas, particularly within the Anatidae family. Moreover, they emphasize the importance of incorporating additional genetic markers, such as nuclear DNA (e.g., microsatellites), to gain a more comprehensive understanding of contemporary migration patterns and population structure in migratory waterfowl.

Given the escalating impact of human activities and climate change on wetland ecosystems, our findings stress the urgent need for regular genetic monitoring to preserve the biodiversity and ecological integrity of these habitats. We recommend conducting genetic assessments every five years to track genetic diversity and population connectivity among migratory species. This proactive approach will provide valuable data to inform conservation strategies, ensuring the long-term survival of migratory birds that rely on the wetlands of the Kashmir Valley during winter months.

In conclusion, this study contributes essential insights into the genetic diversity and phylogenetic relationships of water birds in the Kashmir Himalayas. It highlights the significance of DNA barcoding as a tool for species identification and evolutionary studies, while also advocating for more robust, multilocus genetic analyses to better inform conservation efforts in the face of Anthropocene challenges. Protecting these critical wetland habitats is not only vital for regional biodiversity but also for maintaining the migratory pathways of globally significant species like *Anas crecca*.

## Declarations

### Ethical Approval statement

The collection of samples for this study was conducted during a bird ringing and satellite tagging research program. All procedures adhered to relevant ethical guidelines and regulations and all necessary permissions and licenses were obtained to ensure compliance with legal and ethical standards for wildlife research. The necessary permissions for capture and tagging of birds were provided by the concerned statutory authorities, the Chief Wildlife Warden of Jammu & Kashmir Government and the Ministry of Environment, Forests and Climate Change, Government of India (GOI). Additionally, permits for the import and use of satellite PTTs were issued by the Director General of Foreign Trade, Government of India.

### Consent to Participate (Ethics)

Not Applicable

### Consent for Publication (Ethics)

This study has not used any secondary data for publication, hence no consent to publish was required.

## Availability of data and material

All the DNA sequences generated during the current study are submitted to NCBI (https://www.ncbi.nlm.nih.gov). The accession numbers for each sequences of different genes submitted to NCBI are ON406326–ON406350 for 16S rRNA; for 12S rRNA – ON417475 – ON417521; for COI – ON387719–ON387745; for Dloop–ON400515–ON400539; Cyt b– ON412323–ON412374.

## Author contribution

The first and corresponding author Khursheed Ahmad conceptualized the proposal and designed the framework and acquired funding and resources and prepared the paper manuscript, S.P. Goyal designed framework for genetic analysis and sequencing and helped with paper manuscript, Parag Nigam helped in design and implementation of the project work in field and lab, Divyanshi Bisht and Bhem Dutt Joshi helped in carrying out sequencing and data analysis in the laboratory. Khurshid Alam Khan, Iqram ul Haq and Mohsin Munshi helped in data and sample collection in the field. All the authors read and approved the final version of the manuscript.

## Acknowledgement

we are thankful to Science & Engineering Research Board (SERB) and National Mission for Himalayan Studies (NMHS), Govt. of India for funding this research study. We are highly grateful to the Hon’ble Vice-Chancellor SKUAST-Kashmir, Shalimar and the Chief Wildlife Wardens of Jammu & Kashmir Government and the field and frontline staff of the Department of Wildlife Protection, J&K Government and Director Wildlife Institute of India, for their help and support in successful implementation of this project.

