## Supplementary tables and figures for "Understanding genetic diversity and phylogeography of Common Teal and its phylogenetic relationship with other water bird species in the wetlands of Kashmir Himalayas"

Table S1: Nucleotide Composition of *Anas crecca* with the global data

| Gene | Nucleotide Composition | | | |
| --- | --- | --- | --- | --- |
|  | T% | C% | A% | G% |
| 12sRNA | 16.65 | 29.35 | 30.23 | 23.77 |
| 16sRNA | 21.37 | 26.37 | 30.87 | 21.39 |
| Cyt b | 24.20 | 31.73 | 23.99 | 20.08 |
| Control region | 30.28 | 31.92 | 19.05 | 18.75 |
| COI | 23.71 | 36.55 | 24.05 | 15.69 |

Table S2. Selected primers for different regions of mt DNA genome.

| Primer Name | Sequences (5’–3’) | Size, bp | Final BP size used | References |
| --- | --- | --- | --- | --- |
| 16s RNA_F | CGCCTGTTTATCAAAAACAT | 570 | 440 | Palumbi et al., 1991 |
| 16sRNA_R | CTCCATAGGGTCTTCTCGTCTT |  |  |  |
| 12s RNAF | AAAAAGCTTCAAACTGGGATTAGATACCCCACTAT | 438 | 360 | Kocher et al. 1989 |
| 12s RNAR | TGACTGCAGAGGGTGACGGGCGGTGTGT |  |  |  |
| Cyt b F | CCATCCAACATCTCTGCTTGATGAA | 458 | 370 | Moum and Arnason, 2001 |
| Cyt b R | CCATCCAACATCTCAGCATGATGAAA |  |  |  |
| CR1 F | GTTATTTGGTTATGCATATCGTG | 380 | 380 | Sorenson and Fleischer, 1996 |
| CR1 R | CCTGAAGTAGGAACCAGATG |  |  |  |
| COI F | TTCTCCAACCACAAAGACATTGGCAC | 749 | 570 | Hebert et al., 2003 |
| COI R | TAAACTTCAGGGTGACCAAAAAATCA |  |  |  |

Table (S3): Nucleotide Composition of *Anas crecca* with the same family using COI (574bp)

| **Species** | **A (%)** | **T (%)** | **C (%)** | **G (%)** | **Reference** |
| --- | --- | --- | --- | --- | --- |
| *A. crecca* | 24.05 | 23.71 | 36.54 | 15.69 | Present Study |
| *A. galericulata* | 22.82 | 24.74 | 36.41 | 16.03 | (KJ169568.1) |
| *A. acuta* | 23.52 | 23.69 | 36.41 | 16.38 | (KF312717.1) |
| *A. clypeata* | 23.34 | 26.13 | 33.80 | 16.72 | (KT345702.1) |
| *A. crecca* | 24.22 | 23.69 | 36.41 | 15.68 | (KF203133.1) |
| *A. falcata* | 24.22 | 23.87 | 36.06 | 15.85 | (NC_023352.1) |
| *A. formosa* | 23.52 | 24.22 | 36.06 | 16.20 | (NC_015482.1) |
| *A. platyrhynchos* | 23.52 | 24.39 | 35.89 | 16.20 | (EU009397.1) |
| *A. poecilorhyncha* | 23.52 | 24.39 | 35.89 | 16.20 | (KF156760.1) |
| *A. albifrons* | 24.22 | 25.61 | 34.84 | 15.33 | (NC_004539.1) |
| *A. anser* | 24.22 | 24.91 | 35.37 | 15.51 | (NC_011196.1) |
| *A. cygnoies* | 24.22 | 24.91 | 35.37 | 15.51 | (NC_023832.1) |
| *A. fabalis* | 25.09 | 25.26 | 34.67 | 14.98 | (NC_016922.1) |
| *A. indicus* | 24.04 | 25.26 | 35.19 | 15.51 | (NC_025654.1) |
| *A. semipalmata* | 23.69 | 27.18 | 33.45 | 15.68 | (NC_005933.1) |
| *A. americana* | 23.34 | 23.69 | 36.93 | 16.03 | (NC_000877.) |
| *A. ferina* | 23.52 | 23.69 | 36.93 | 15.85 | (KJ710708.) |
| *A. fuligul* | 23.34 | 24.39 | 36.24 | 16.03 | (KJ722069.1) |
| *B. bernicla* | 23.87 | 25.44 | 34.49 | 16.20 | (KJ680301.1) |
| *B. canadensis* | 23.34 | 23.87 | 36.41 | 16.38 | (NC_007011.1) |
| *C. moschata* | 22.30 | 24.39 | 36.41 | 16.90 | (NC_012843.1) |
| *C. atratus* | 23.34 | 24.56 | 35.37 | 16.72 | (NC_012843.1) |
| *C. columbianus* | 24.56 | 24.91 | 34.84 | 15.68 | (NC_007691.1) |
| *C. cygnus* | 24.22 | 25.09 | 34.84 | 15.85 | (NC_027095.1) |
| *C. olor* | 24.04 | 24.91 | 34.67 | 16.38 | (NC_027096.1) |
| *D. javanica* | 25.09 | 26.66 | 31.36 | 16.90 | (NC_012844.1) |
| *M. merganser* | 23.69 | 23.87 | 36.41 | 16.03 | (NC_016723.1) |
| *M. squamatus* | 23.17 | 23.87 | 36.41 | 16.55 | (NC_016723.1_) |
| *N. rufina* | 24.56 | 25.44 | 35.02 | 14.98 | (NC_024922.1_) |
| *T. ferruginea* | 22.30 | 24.39 | 36.06 | 17.25 | (NC_024640.1) |
| *T. tadorna* | 23.34 | 23.87 | 36.41 | 16.38 | (KU140668.1) |

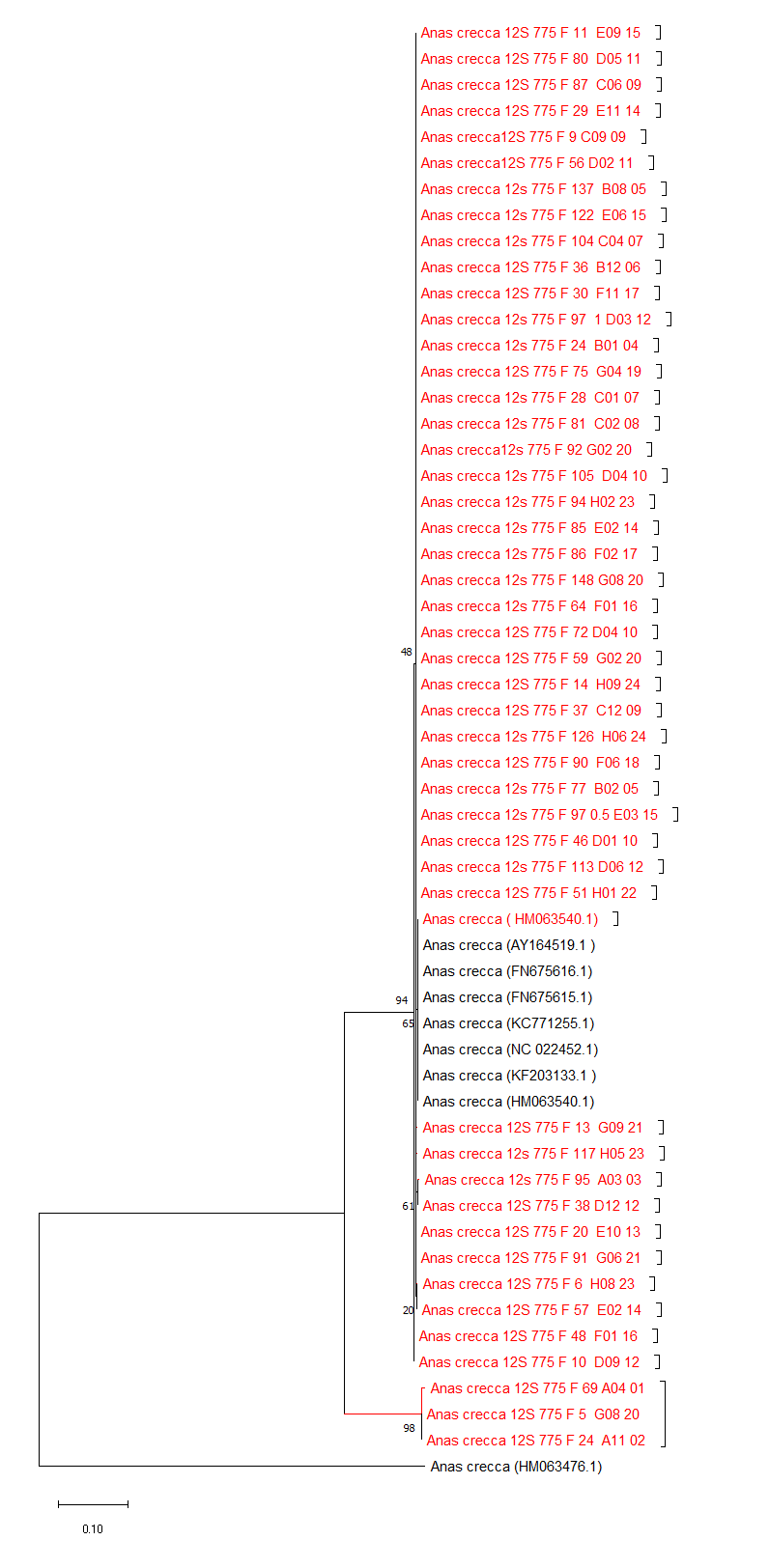

Figure S1:- Maximum Likelihood Tree of 12s gene of *Anas crecca* with the global data. (Red color indicates sequences generated in present study; numbers in nodes are bootstraps values).

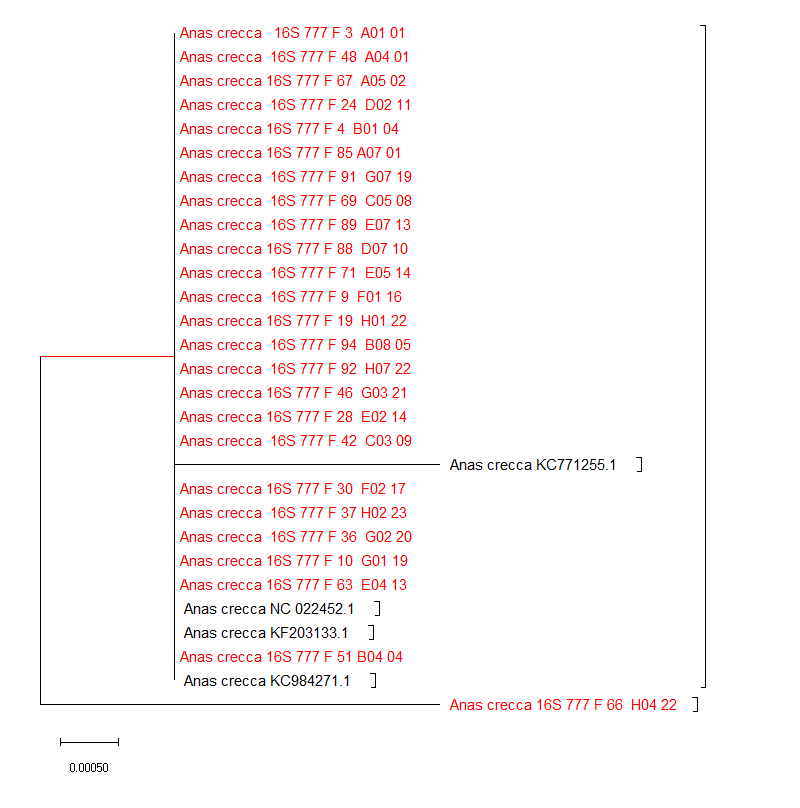

Figure S2:- Maximum Likelihood Tree of 16s gene of *Anas crecca* with the global data. (Red color indicates sequences generated in present study; numbers in nodes are bootstraps values).

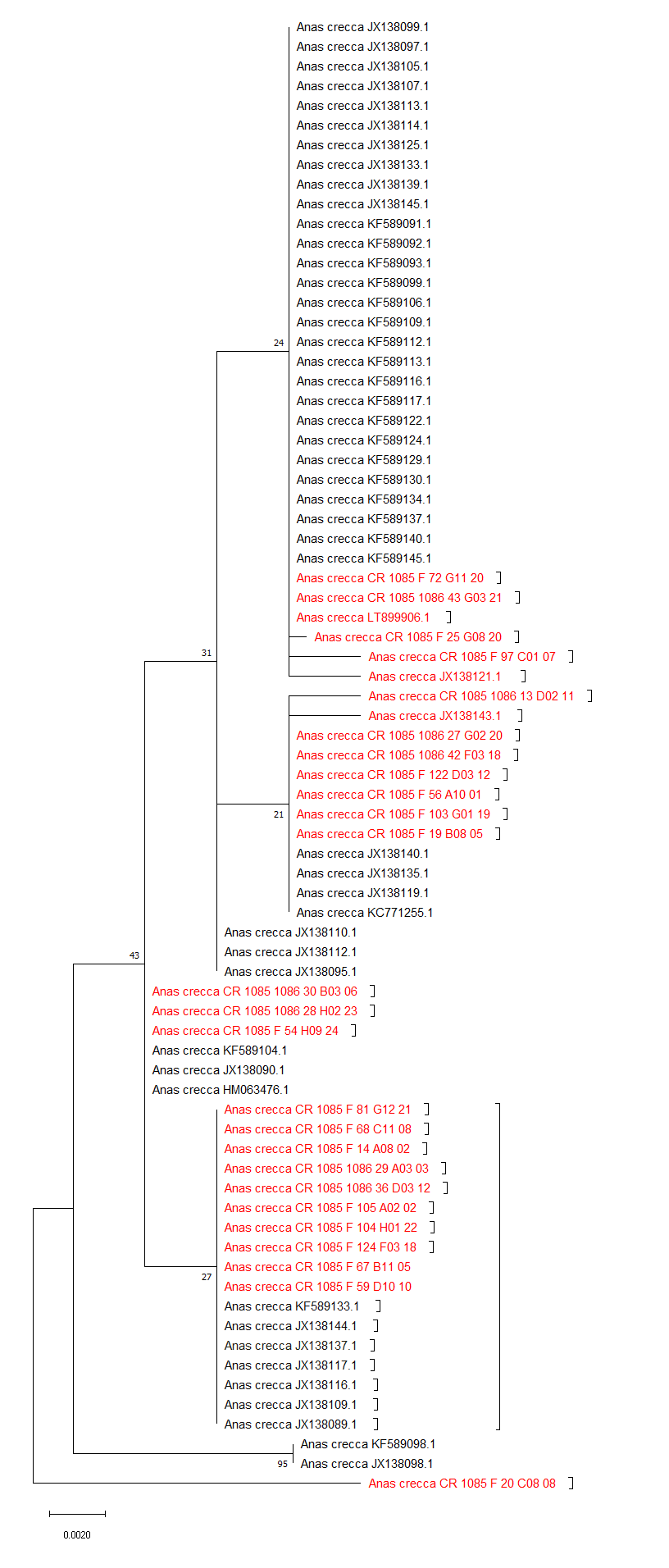

Figure S 3:- Maximum Likelihood Tree of CR gene of *Anas crecca* with the global data. (Red color indicates sequences generated in present study; numbers in nodes are bootstraps values).

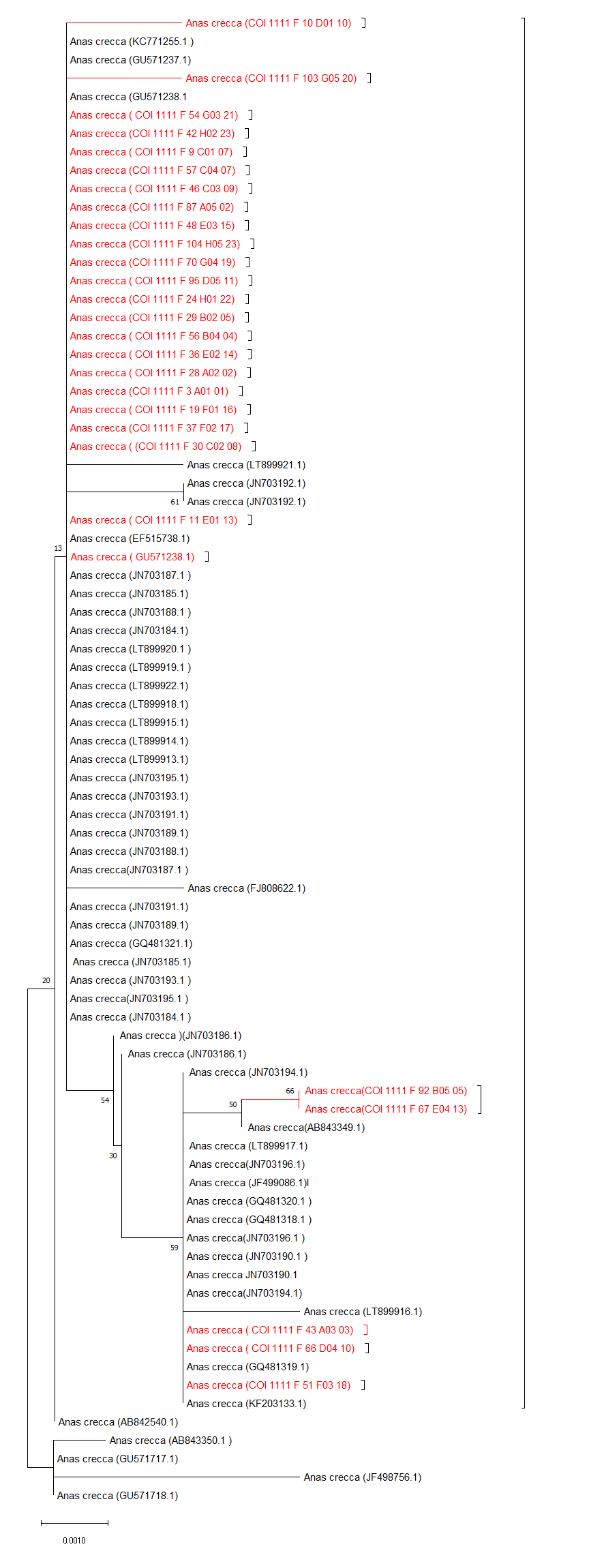

Figure S4:- Maximum Likelihood Tree of COI gene of *Anas crecca* with the global data. (Red color indicates sequences generated in present study; numbers in nodes are bootstraps values).

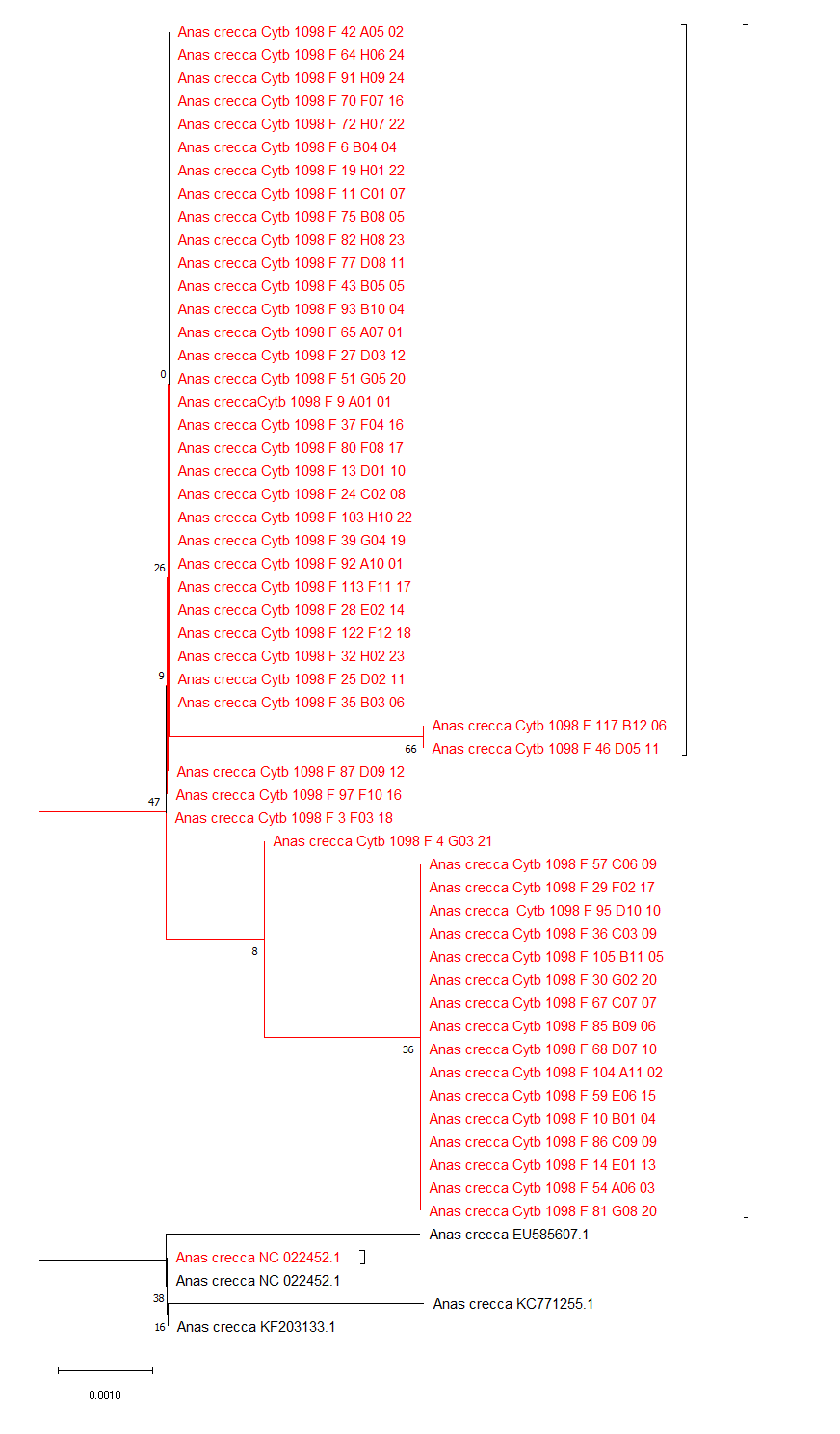

Figure S5:- Maximum Likelihood Tree of Cytb gene of *Anas crecca* with the global data. (Red color indicates sequences generated in present study; numbers in nodes are bootstraps values).

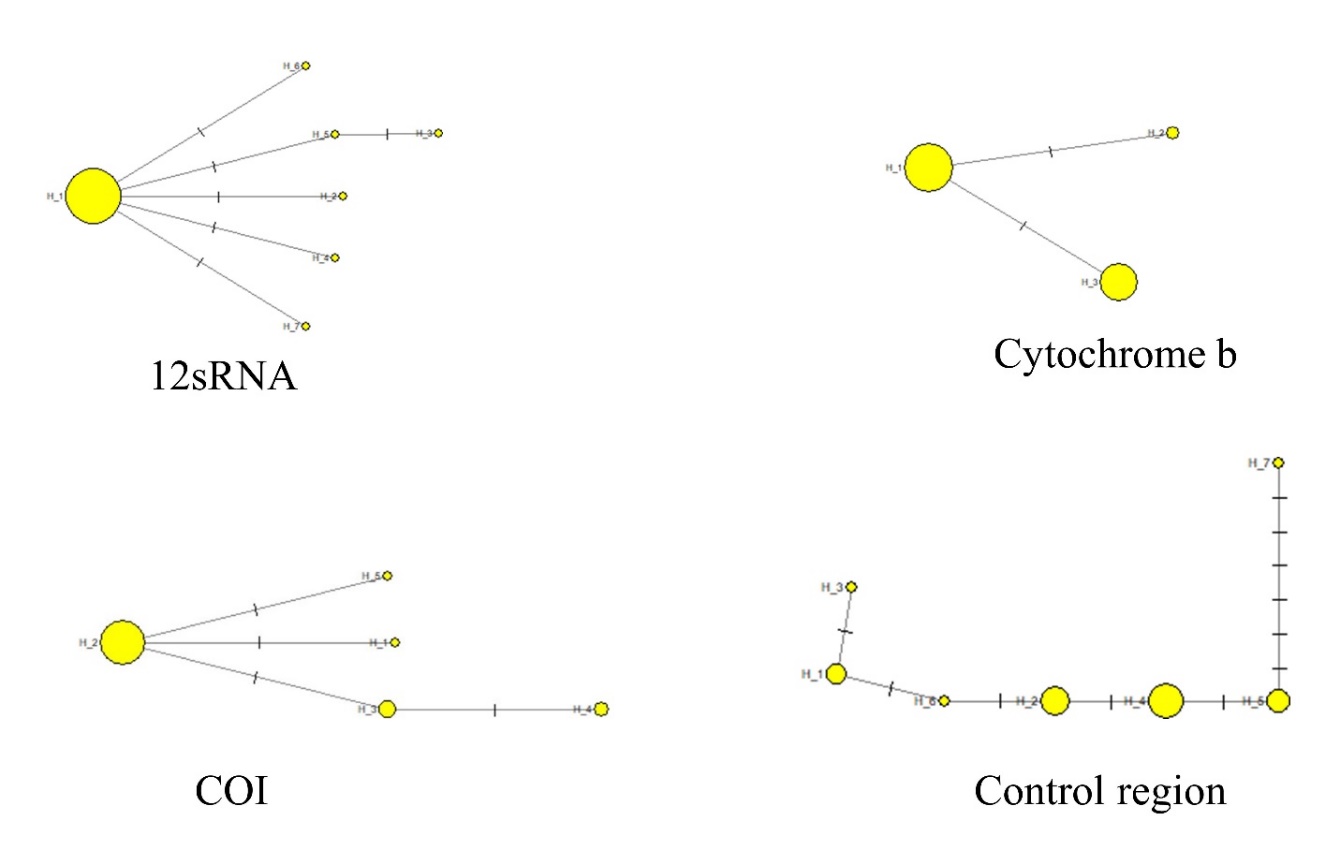

Figure S6. Minimum spanning network of observed haplotypes in Common Teal (*Anas crecca*) based on different mtDNA genes. Circle size is in proportional number of individuals.
